# Genetic Diversity of Non-Tuberculous Mycobacteria among Symptomatic TB negative Patients in Kenya

**DOI:** 10.1101/2021.10.28.465104

**Authors:** Zakayo M Mwangi, Nellie Mukiri, Frank G Onyambu, Wallace Bulimo

## Abstract

Non-Tuberculous Mycobacteria (NTM) transmission to humans occurs through inhalation of dust particles or vaporized water containing NTM leading to pulmonary manifestations. NTM infections are often misdiagnosed for tuberculosis (TB) due to their similar clinical and radiological manifestations. We, therefore, performed a species-level identification of NTM in symptomatic TB negative patients through sequencing of the *hsp65* gene. We conducted a cross-sectional study at the National Tuberculosis Reference Laboratory in the period between January to November 2020. One hundred and sixty-six mycobacterial culture-positive samples that tested negative for TB using capilia underwent Polymerase Chain Reaction targeting the *hsp65* gene. Isolates showing a band with gel electrophoresis at 441 bp position were sequenced using Sanger technology. Geneious software was used to analyze the obtained sequences, and the National Center for Biotechnology Information gene database identified NTM species for each isolate. A phylogenetic tree was constructed from the DNA sequences and evolutionary distances computed using the general time-reversible method. Pearson chi-square was used to determine the association between NTM infection and participants’ characteristics

Our study identified 43 different NTM species. The dominant NTM belonged to *Mycobacterium avium complex* 37 (31%). Slow-growing NTM were the majority at 86 (71%) while rapid-growing NTM were 36 (29%). A significant association (p<0.05) was observed for regions and age, while patient type had a week likelihood of NTM infection. Our study characterized the diversity of NTM in Kenya for the first time and showed a high diversity of NTM species.

## Introduction

NTM are ubiquitous environmental organisms that are present in habitats such as water, soil, and dust. While in the environment, NTM colonies coalesce in clusters forming biofilms that enable their persistence in adverse conditions such as harsh weather and degradation by disinfectants[1]. The biofilms’ hydrophobicity enhances NTM dispersion and subsequent transmission to humans mainly through inhalation of dust particles or vaporized water containing NTM leading to NTM pulmonary disease (NTM-PD)[2], [3], but there is no evidence of human to human transmission [4]. Environmental and climatic conditions influence the distribution of more than the 200 NTM species [5] with specific species predominating various geographic regions [6], [7]. For instance, *Mycobacterium avium complex (MAC), M. gordonae* and *M. xenopi* were dominant in the different geographical regions within Europe [8], in the Middle East *M. abscessus, M. fortuitum*, and *M. intracellulare* were commonly isolated [9], *M*.*fortuitum* was the most common NTM in India [10]. MAC was dominant in Singapore and Japan [10] while *MAC, M*.*intracellulare*, and *M*.*kansasii* were the most common in Sub-Saharan Africa [11].

The NTM-PD clinical presentation is similar to that of *Mycobacteria tuberculosis* (MTB) and is often misdiagnosed for MTB. This leads to treatment complications since MTB and NTM management strategies are incongruent [12], [13]. Identification of NTM is important in the determination of the clinically relevant species and deciding on the appropriate treatment regimen for the infection since mycobacterium susceptibility to antimicrobial agents varies with individual species [13]–[16]. Classical NTM identification and species differentiation procedures are simple and cost-effective and utilize phenotypic characteristics such as their growth rate, pigment production, growth at different temperatures, and biochemical analysis including niacin accumulation test, nitrate reduction test, urea hydrolysis among several others [17]–[19]. These methods however have limitations in NTM diagnosis due to their time-consuming, tedious nature, and lack of accurate distinction of closely related species [17], [20].

Molecular methods are seen to be sufficient and highly efficient in the discrimination of NTM species in a fast and reliable manner a property that phenotypic methods lack [5], [21], [22]. The commonly applied molecular technique is by targeted sequencing of common housekeeping genes including *16s rRNA, rpoB*, and *hsp65*. The sequences of these genes exhibit immense discriminatory power in their identification of the NTM species, hence providing a robust phylogenetic tree that enables a proper classification of NTM to either rapid or slow growers, and gives a better understanding of their evolutionary diversity[23]. Further, concatenation of the sequences belonging to these genes provides for the description and characterization of novel NTM, hence increasing the number of identifiable NTM species in the gene databases [24]–[26]. Line probe assays (LPA) is another molecular technique that is gaining popularity, especially in microbiology reference laboratories in resource-limited settings [27], [28]. This assay employs the reverse hybridization technique where amplicons are hybridized on a nitrocellulose membrane strip and result interpretation is done based on the presence of bands at various points on the strip enabling for the simultaneous detection and identification of NTM species[17], [29]. The commonly used LPA is GenoType® *Mycobacterium* CM/AS assay (*Hain* Lifescience, Nehren Germany) which is a commercial kit that relies on partial amplification of the 23S rRNA.

The 16s rRNA is composed of highly conserved as well as variable regions making it the commonly studied gene for bacterial identification and a resourceful target for studying phylogenetic relationships [20]. This gene however shows high inter-species sequence similarity with some mycobacteria having absolute homology in their sequences. For instance, *M. abscessus/M. bolletii/M. massiliense, M. austroafricanum/M. vanbaalenii, M. kansasii/M. gastri, M. senegalense/M. houstonense and, M. mucogenicum/M. phocaicum* present with 0% inter-species divergence making the 16s rRNA gene less suitable for mycobacteria species differentiation [30]. The *rpoB* gene encodes for the *β* subunit of the RNA polymerase an enzyme responsible for RNA synthesis [31]. This gene has a higher sensitivity in mycobacteria species differentiation with an inter-species similarity score ranging from 92.71% to 100% compared to that of 16s rRNA at 96.57% to 100% [32]–[34]. The *hsp65* gene belongs to the heat shock protein family that is involved in intracellular protein folding, assembly, and transport thus being highly immunogenic. Compared to 16s rRNA and *rpoB*, the *hsp65* gene sequence is more diverse with a high genetic heterogeneity and a lower inter-species similarity percentage ranging from 89.2% to 100 %,therefore producing a more robust phylogenetic tree with most nodes having bootstrap value above 80% [17], [23], [32], [35], [36].

The application of the above-mentioned molecular diagnostic techniques that detect and distinguish NTM species have played a major role in the increased reporting of NTM cases especially in Sub-Saharan Africa [11]. Detection of NTM in Kenya has relied on LPA which is limited to identifying NTM complexes with the GenoType Mycobacteria CM/AS recognizing 15 and 16 NTM species respectively, hence limiting a comprehensive overview on NTM species diversity [17], [27].

In this study we obtained sputum samples from symptomatic TB culture-negative patients and identified the NTM species through sequencing of the *hsp65* gene, and also described their evolutionary diversity. NTM species obtained by *hsp65* sequencing were also compared to those obtained using line probe assay (GenoType Mycobacteria CM/AS).

## Materials and methods

### Samples

Sputum samples received at the National Tuberculosis Reference Laboratory (NTRL) underwent mycobacterial culture and identification procedures. A total of 165 NTM samples received from January to November 2020 were aliquoted and transferred to the Molecular and Infectious Diseases Research Laboratory (MIDR-L) for sub-culturing and *hsp65* PCR.

### Laboratory procedures

#### Sample processing, mycobacterial culture, and growth identification

The sputum samples were decontaminated using the N-acetyl-L-cysteine 2% NaOH (NALC-NaOH) procedure, then inoculated into Mycobacteria Growth Indicator Tube (MGIT) and Lowenstein-Jensen (LJ) media, incubated at 37°C and monitored for growth for up to 6 and 8 weeks respectively. At the same time, sputum smears were prepared, air dried, heat-fixed then fluorochrome stained with auramine O where mycobacteria appeared as bright yellow fluorescent rods when viewed under a light-emitting diodes (LED) microscope.

The culture growth in MGIT and LJ underwent the Mtb identification testing using the SD Bioline TB Ag MPT64 assay (capilia) (Standard Diagnostics, Yongin-si, Gyeonggi-do, Republic of Korea), and capilia positive samples were excluded from the study. The capilia negative samples underwent ZN microscopy with the presence of AFB indicating a possible NTM.

### DNA extraction

Mycobacterial DNA was extracted from 500 μL of re-suspended colonies using GenoLyse^®^ (*Hain* Lifescience, Nehren, Germany) according to the manufacturer’s instructions. Briefly, 100ul of lysis buffer (A-LYS) buffer was added to each cryovial containing the resuspended colonies and incubated for 5 minutes at 95°C after which 100ul neutralization buffer (A-NB) was added and centrifugation was done at 5000G for 10 minutes. The supernatant was transferred to a newly labeled cryovial awaiting PCR.

### Conventional PCR, gel electrophoresis, and DNA purification

A conventional PCR targeting the *hsp65* gene was conducted using the GoTaq® Green Master Mix (Promega, Madison, Wisconsin, USA) in a final reaction volume of 13μl comprising 6.25μl of 2X GoTaq Hot Start Green Master Mix, 2.5μl DNA template, 0.25μl of each of both F- (5′-ACCAACGATGGTGTGTCCAT-3′) and R- (5′-CTTGTCGAACCGCATACCCT-3′) primers at a final concentration of 10 pmoles, and 3.75μl of nuclease-free water to make up the reaction volume. Thermal cycling conditions were 1 cycle of 94°C for 4 minutes, 35 cycles of 94°C for 1 minute, 57°C for 1 minute, 72°C for 1 minute and a final extension for 10 minutes at 72°C. Amplified products were confirmed on a 1% Agarose gel stained with 4.6μl SYBR safe DNA stain (Invitrogen, Carlsbad, California, USA), and bands at 441 bp were observed in an Ultra Violet gel viewer. The PCR products were enzymatically purified using ExoSAP IT (Applied Biosystems, Foster City, California, USA). Purification conditions were 37ºc for 15 minutes followed by a second incubation at 80ºc for 15 minutes and a final cooling step at 4ºC for 5 minutes.

#### *hsp65* sequencing

The purified amplicons were sequenced in the forward direction by Sanger sequencing using Big Dye™ Terminator Version 3.1 Cycle Sequencing Kit (Applied Biosystems, Foster City, California, USA) and the forward primer. The sequencing reaction was a 10μl reaction comprising 1.25 μl of Big Dye Terminator, 3 μl of 5X Sequencing Buffer, 1 μl of 1 pmol of the sequencing primer, and 1.5 μl of the PCR product. The reaction volume was made up by adding 3.25 μl of nuclease-free water. The reaction proceeded through 96°C for 1 minute then 25 cycles of 96°C for 10 seconds, 50°C for 5 seconds, and 60°C for 4 minutes.

Purification of cycle-sequencing products was done using the BigDye X Terminator™ purification kit following manufacturer’s instructions (Applied Biosystems, Foster City, California, USA) and purified products were loaded onto the ABI 3730 genetic analyzer (Applied Biosystems, Foster City, California, USA) for capillary electrophoresis.

### Data analysis

Geneious software was used to align raw AB1 files from the ABI 3730 then sequences for each sample were compared with those in the GenBank (National Center for Biotechnology Information: http://www.ncbi.nlm.nih.gov/) and (*hsp65*)-BLAST (hsp65-BLAST) (http://hsp65blast.phsa.ca/blast/blast.html) DNA sequence database. NTM species identification was confirmed if a 97% match was achieved.

Statistical data analyses were performed using STATA version 13 (Statacorp LLC, Texas, USA). The Pearson’s chi-square *χ*2 test was used to compare differences in proportions; variables with *P*< 0.05 were considered statistically significant.

### Phylogenetic analysis

Sequences of the *hsp65* were trimmed to start and end at the same nucleotide position for all isolates. The alignment of multiple sequences was conducted with Geneious prime 2020.2 software then MEGA X version 10.2.4 software performed phylogenetic analysis on the 440bp sequence.

The phylogenetic tree was constructed from the DNA sequences by using the neighbor-joining method, and the evolutionary distances were computed using the general time-reversible method. The gene sequence for *M. tuberculosis* H37Rv was used as the phylogenetic root.

## Results

Polymerase chain reaction of the *hsp65* gene was performed to all the 166 samples and a DNA band at approximately 441bp was observed with agarose gel for 146 (88.5%) isolates. Twenty (11.5%) isolates did not show a band during gel electrophoresis (C9, C22, C37, C38, C45, C50, C51, C55, C63, C73, C85, C97, C105, C112, C113, C114, C120, C135, C149, and C161) and were excluded from further analysis. Sequence analysis of the *hsp65* for the 146 PCR products showed NTM species in 122 (84%) of the isolates.

A total of 50 different species were identified with NTM comprising 43 (86%), and 7(14%) were found to be organisms other than NTM. The non-NTM species observed include *M. tuberculosis, Rothia spp, Norcadia spp, Kocuria spp, Streptomyces spp, Rhodococcus spp*, and *Planctomycetes spp*

The frequently isolated NTM belonged to *Mycobacterium avium complex* (31%) (*M. avium* subspecies, *M. intracellulare, M. colombiense, M. yongonese, M. parascrofulaceum, M. paratuberculosis*, and *M. timonense*). Second most isolated NTM species belonged to *M. fortuitum complex* (20%) (*M. fortuitum subsp fortuitum, M. conceptionense, M. senegalense, M. peregrinum, M. mageritense, M. neworleansense, and M. porcinum*), followed by *M. abscessus complex species* (14%) (*M. abscessus subsp. bollettii and M. abscessus subsp. abscessus*). The species belonging to these three complexes accounted for 65% of all mycobacteria identified.

Most samples came from patients living in the Coastal region (41%), followed by Nairobi (26%) region. The Coastal region had the highest (18%) variety of NTM species with *M. fortuitum* (44%) being the most common NTM. The median age of participants was 39 years (Interquartile range, IQR 20–50) with the majority belonging to the age group between 30-39 years. The majority of subjects with NTM were males (73%). Patients on treatment follow-up for multi-drug resistant tuberculosis (MDR-TB) (31%) were the majority (Table 1).

**Table 1:** Sociodemographic characteristics of patients with NTM.

| Variables | Participants n (%) | participants with NTM | X<sup>2</sup>, p value |
| --- | --- | --- | --- |
| **Regions** |  |  |  |
| Mt. Kenya | 27(16) | 25(19) | 402, 0.002* |
| North Rift Valley | 22(13) | 13(10) |  |
| South Rift valley | 22(13) | 12(8) |  |
| Lake Victoria | 15(9) | 8(5) |  |
| Lower Eastern | 9(5) | 8(5) |  |
| Coast | 41(25) | 37(30) |  |
| North eastern | 4(2) | 4(3) |  |
| Nairobi | 26(16) | 28(20) |  |
| Subtotal | 166(100) | 122(100) |  |
| **Age in years** |  |  |  |
| <21 | 7(4) | 5(4) | 0.000* |
| 21-29 | 36(22) | 26(21) |  |
| 30-39 | 44(27) | 35(28) |  |
| 40-49 | 37(22) | 25(19) |  |
| 50-59 | 23(14) | 15(14) |  |
| 60-69 | 11(7) | 10(9) |  |
| 70-79 | 6(4) | 5(4) |  |
| >79 | 2(1) | 1(1) |  |
| Subtotal | 166(100) | 122(100) |  |
| **Gender** |  |  |  |
| Female | 45(27) | 32 (27) | 57, 0.134 |
| male | 121(73) | 90(73) |  |
| Subtotal | 166(100) | 122(100) |  |
| **Type of patient** |  |  |  |
| new | 47(28) | 33(27) | 213, 0.074 |
| retreatment | 28(17) | 21(17) |  |
| relapse | 39(23) | 30(25) |  |
| MDR follow-up | 52(31) | 38(31) |  |
| Subtotal | 166(100) | 122(100) |  |

To investigate the genetic relatedness of the isolated NTM, a phylogenetic tree was constructed. Phylogenetic analysis based on the *hsp65* genes revealed very close genetic similarity in the NTM species, with clear separation of the slowly and rapidly growing mycobacteria. Slow growing NTM were the majority at 86 (71%) (Table 2) while rapid-growing NTM were 36 (29%) (Table 3). Each mycobacterial species was identified as a distinct entity in the phylogenetic tree (Figure 1).

**Table 2:**
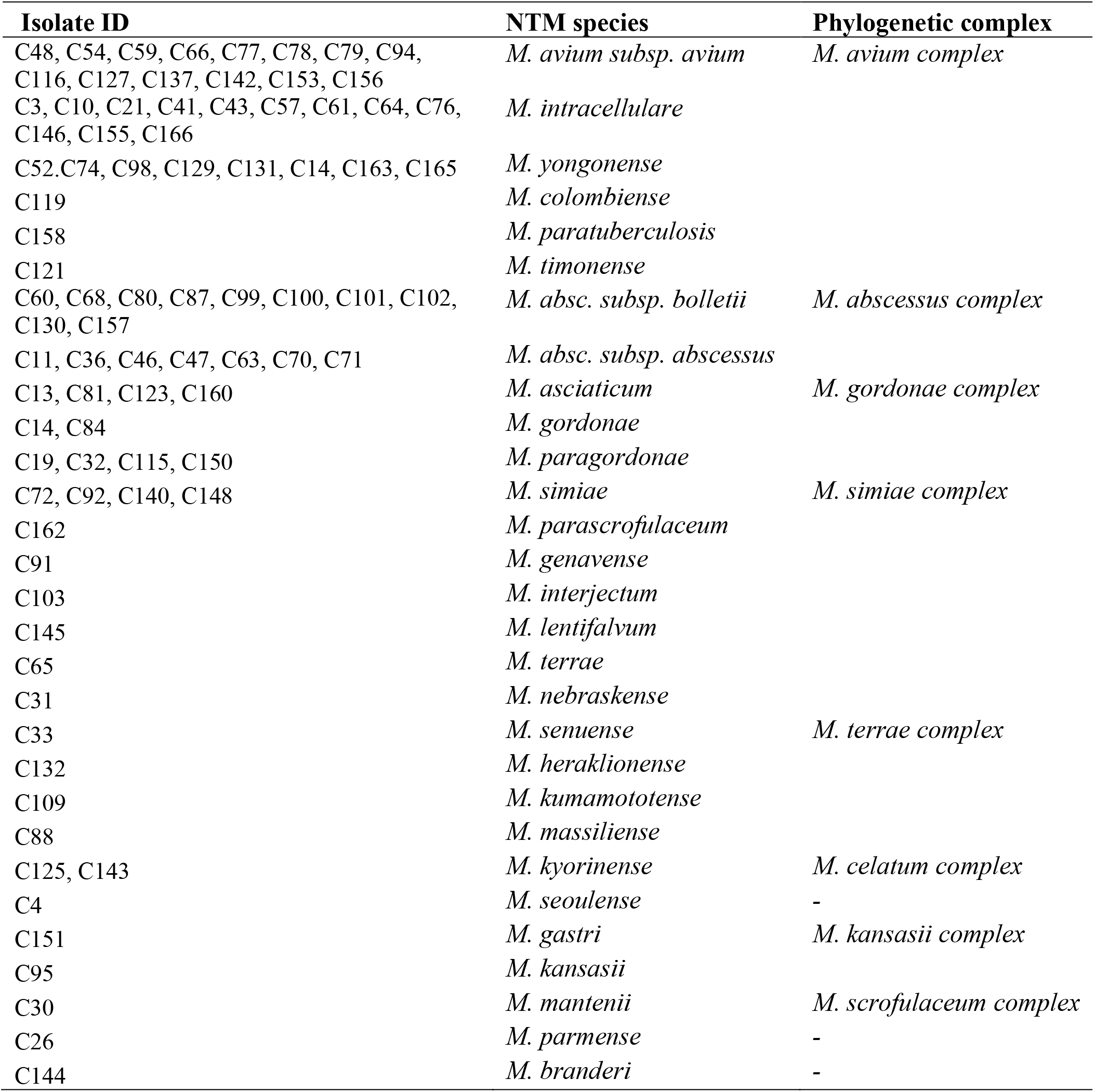
Frequency of slow-growing NTM and their phylogenetic complexes identified by *hsp65* partial sequencing.

**Table 3:** Frequency of rapidly growing NTM and their phylogenetic complexes identified by *hsp65* partial sequencing.

| **Isolate ID** | **NTM species** | **Phylogenetic complex** |
| --- | --- | --- |
| C16, C35, C40, C67, C96, C108, C124, C126, C128, C134 | *M. fort. subsp fortuitum* | *M. fortuitum* complex |
| C49, C90, C107, C152 | *M. conceptionense* |  |
| C28, C106, C118, C147 | *M. senegalense* |  |
| C110, C159, C164 | *M. peregrinum* |  |
| C24 | *M. mageritense* |  |
| C122 | *M. neworleansense* |  |
| C27 | *M. porcinum* |  |
| C1, C15C39, C93 | *M. monacense* | - |
| C12, C42, C53 | *M. chelonae* | *M. chelonae* complex |
| C104 | *M. phocaicum* | *M. mucogenicum* complex |
| C6 | *M. neoaurum* | *M. parafortuitum* complex |
| C18 | *M. manitobense* | - |
| C58 | *M. alsense* | - |
| C133 | *M. bourgelatii* | - |

**Fig. 1.**
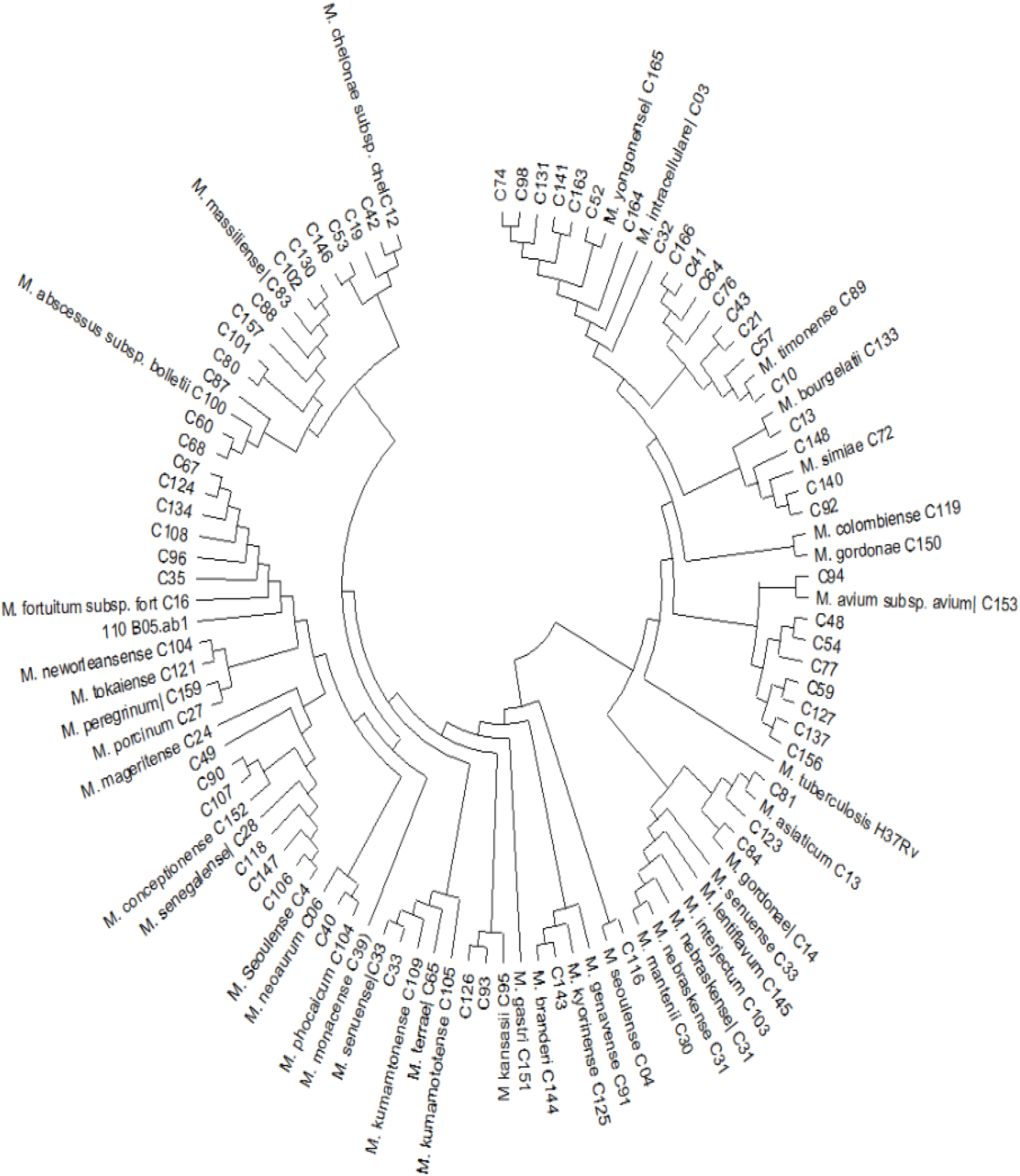
Phylogenetic tree showing the diversity of NTM species isolated from sputum samples in symptomatic TB negative patients in Kenya. The tree was rooted to Mycobacterium tuberculosis H37Rv which was also used as a positive control in the analysis.

The results of NTM identified through *hsp65* gene sequencing and GenoType Mycobacterium CM/AS (Hain Lifesciences, Nehren, Germany) were compared for the 122 isolates. Eighty-two (67%) were identical in speciation of NTM (Table 4) while 24 (19%) were discordant for both assays (Table 5). The GenoType Mycobacterium CM/AS assay was unable to identify 16 (13%) NTM species that were later on identified by *hsp65* gene sequencing (Table 6).

**Table 4:**
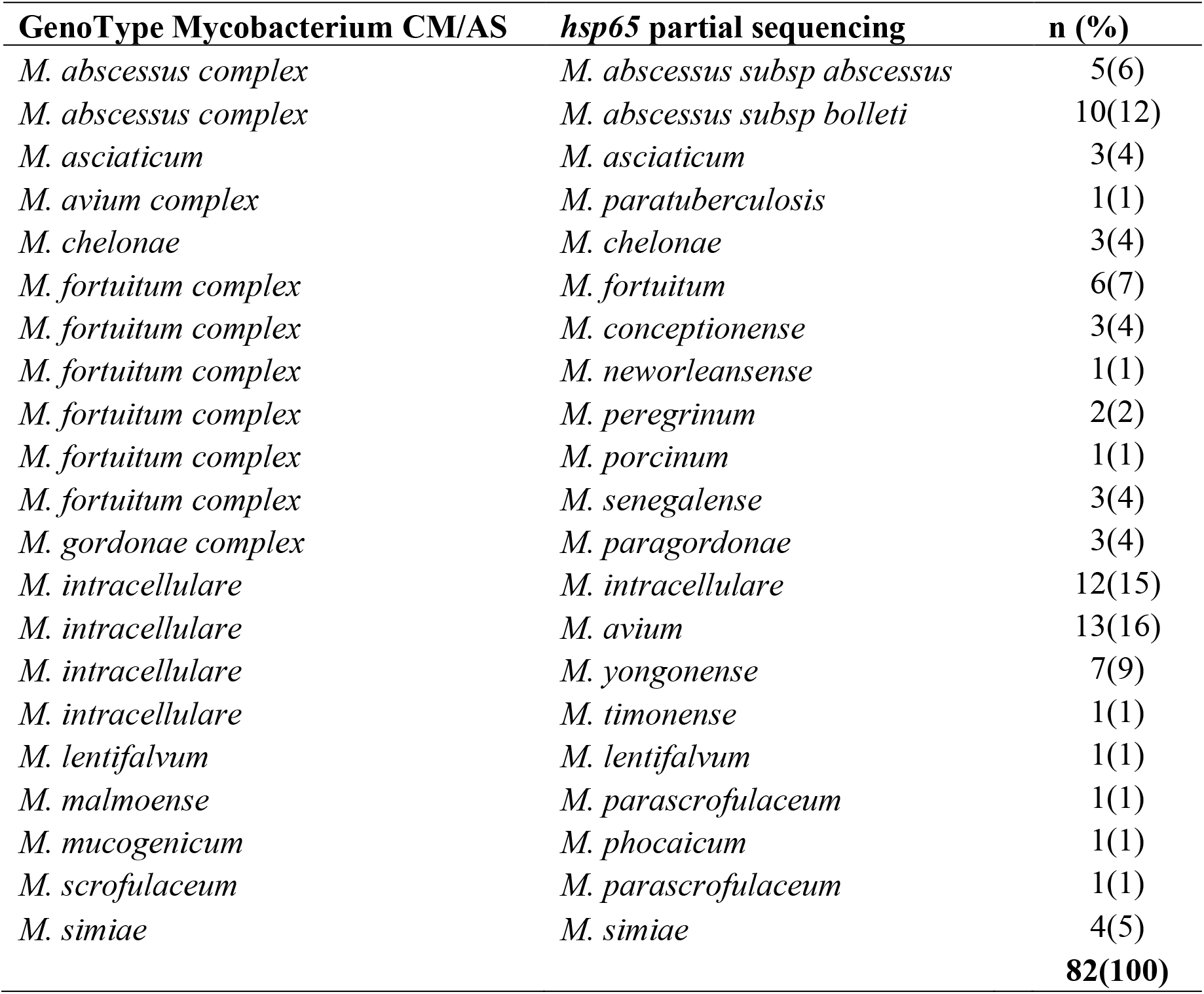
Similar results for NTM speciation by Genotype Mycobacterium CM/AS and *hsp65* gene sequencing.

**Table 5:**
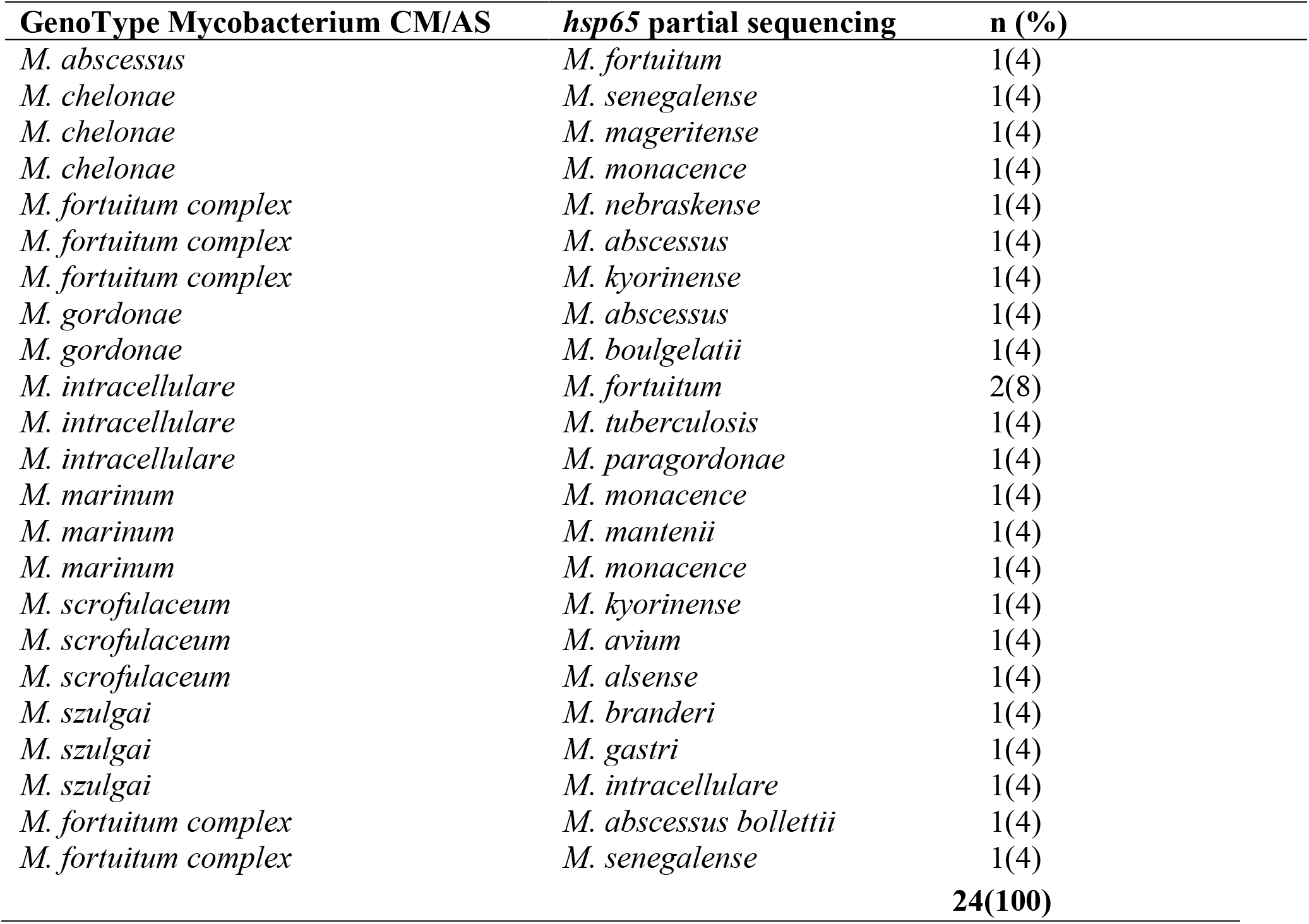
Discordant results in NTM identification by Genotype Mycobacterium CM/AS and *hsp65* partial sequencing.

| **GenoType Mycobacterium CM/AS** | ***hsp65* partial sequencing** | **n (%)** |
| --- | --- | --- |
| *M. abscessus* | *M. fortuitum* | 1(4) |
| *M. chelonae* | *M. senegalense* | 1(4) |
| *M. chelonae* | *M. mageritense* | 1(4) |
| *M. chelonae* | *M. monacence* | 1(4) |
| *M. fortuitum complex* | *M. nebraskense* | 1(4) |
| *M. fortuitum complex* | *M. abscessus* | 1(4) |
| *M. fortuitum complex* | *M. kyorinense* | 1(4) |
| *M. gordonae* | *M. abscessus* | 1(4) |
| *M. gordonae* | *M. boulgelatii* | 1(4) |
| *M. intracellulare* | *M. fortuitum* | 2(8) |
| *M. intracellulare* | *M. tuberculosis* | 1(4) |
| *M. intracellulare* | *M. paragordonae* | 1(4) |
| *M. marinum* | *M. monacence* | 1(4) |
| *M. marinum* | *M. mantenii* | 1(4) |
| *M. marinum* | *M. monacence* | 1(4) |
| *M. scrofulaceum* | *M. kyorinense* | 1(4) |
| *M. scrofulaceum* | *M. avium* | 1(4) |
| *M. scrofulaceum* | *M. alsense* | 1(4) |
| *M. szulgai* | *M. branderi* | 1(4) |
| *M. szulgai* | *M. gastri* | 1(4) |
| *M. szulgai* | *M. intracellulare* | 1(4) |
| *M. fortuitum complex* | *M. abscessus bollettii* | 1(4) |
| *M. fortuitum complex* | *M. senegalense* | 1(4) |
|  |  | **24(100)** |

**Table 6:**
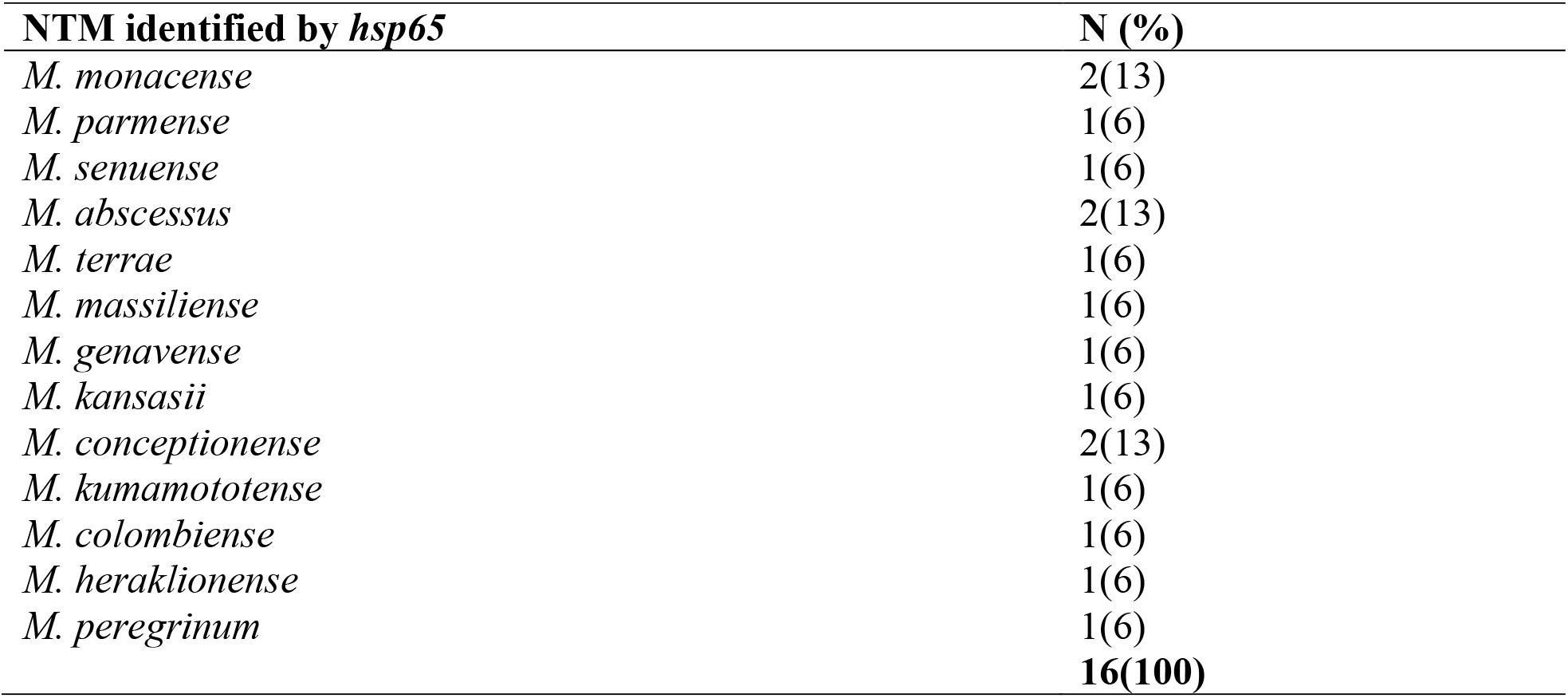
NTM identified by *hsp65* but not identifiable by Genotype Mycobacteria CM/AS.

## DISCUSSION

Global estimates show that MAC is the dominant NTM comprising between 34% and 61% of NTM isolated across the different continents [37], [38]. These findings are consistent with our study that identified species belonging to *MAC* (31%) (*M. avium* subspecies (32%), *M. intracellulare* (27%), *M. yongonese* (18%), *M. parascrofulaceum* (5%), *M. paratuberculosis* (2%), and *M*.*timonense* (2%)) as the most widely isolated NTM in Kenya. Other countries with similar findings include China [39]–[41], Russia [42], America [43], [44], various countries in Europe [45] and Africa [11], [20], [46]–[48]. The high infectivity rate of MAC species could be attributed to their seemingly abundant nature and distribution in different environmental sources such as water and soil, consequently increasing its ease of spread and infection to humans [20], [49]. MAC has been highly associated with NTM-PD [50]; however, it remains unclear how its infectivity relates to NTM-PD [51].

NTM species distribution varies in different geographical locations with environmental conditions contributing towards their distribution patterns [8]. Our findings suggest a regional variation in the diversity of NTM and a possible influence by the different geographical or environmental landscapes within Kenya [11], [36]. For instance, Nairobi is warm and wet with *M*.*senegalense* (15%) being the dominant NTM, Coastal region is very hot and very wet with a majority of the isolates having *M. fortuitum* (44%), *M. abscessus bolletii* (19%) was the most common in the Mt.Kenya region which is cold and wet, North Rift Valley is hot and dry and had *M*.*yongonense* (33%) as the prevalent NTM, and the Western region is hot and wet with *M*.*avium* (33%) as the most common NTM species. These findings may be inconclusive due to the limited number of samples analyzed in our study, and a need for larger and more systematic studies capturing both human and environmental information is required for a more comprehensive understanding of environmental NTM species distribution [8]. These differences in species distribution may partly determine the frequency and manifestations of pulmonary NTM disease in each geographical location.

Certain demographic characteristics such as age and gender have been shown to increase the likelihood of NTM infection [52]. In our study, persons with pulmonary NTM infection were mostly in the youthful age group (30-39 years) and these findings are similar to those of other resource-limited countries [11]. NTM was less likely to infect females suggesting a likelihood of estrogen conferring a protective effect by up-regulating immunological response to clearing of bacterial infections [53], [54].

Pulmonary NTM infection was found to be commonly associated with patients on follow-up for MDR-TB (31%). MDR-TB patients in Zambia and Nigeria recorded similar findings with NTM prevalence of 30% and 56% respectively [52]. Sub-Saharan African countries have the greatest burden of TB infections [55]. Pulmonary TB infections compromise the structural integrity of the lungs subsequently providing a suitable niche for NTM adherence and multiplication, hence the high number of NTM observed in MDR-TB cases [56]. A co-infection of TB and NTM could also be a possible occurrence in MTB-DR cases where the presence of NTM is attributed to its failure to respond to the first-line anti-TB drugs [57].

The partial sequencing of the *hsp65* gene was seen to be more sensitive in its ability to discriminate NTM to the species and subspecies level compared to GenoType Mycobacterium CM/AS (Hain Lifesciences GmbH, Nehren, Germany) [58]. A worthwhile observation was that 10 (71%) of the 14 *M*.*avium* detected by *hsp65* had been identified as *M*.*intracellulare* using Genotype Mycobacterium CM/AS (Hain Lifesciences, Nehren, Germany) suggesting a limited intraspecies detection heterogeneity by Genotype Mycobacterium CM/AS [59]. These occurrences for the Genotype Mycobacterium CM/AS could be attributed to its target gene the 23S rRNA having more conserved and few hypervariable regions, hence limiting its capacity to differentiate closely related NTM [59]. The contrasting results observed between *hsp65* sequencing and Mycobacterium CM/AS in NTM species identification could either be due to mislabeling of sequences in the genetic databases, or a possible co-infection of two different NTM[59].

## Conclusion

Our study characterized the diversity of NTM in Kenya for the first time and identified 43 different NTM species. MAC was the most prevalent in the country followed by *M. fortuitum complex* and *M. abcessus complex* species.

Slow-growing NTM were the majority at 86 (71%) while rapid-growing NTM were 36 (29%). A significant association (p<0.05) was observed for regions and age, while patient type had a week likelihood of NTM infection.

## Limitation

Our study lacked clinical details on the participants, such as underlying lung conditions and HIV status, making it difficult to identify clinically relevant NTM and their association with NTM-PD. Despite these limitations, our study characterized the diversity of NTM in Kenya for the first time and identified that species from the *Mycobacterium avium complex* are the most prevalent in the country. Furthermore, we have shown that the diversity of mycobacteria species is high in Kenya, implying that before treatment of suspected tuberculosis patients, doctors should consider deeper characterization beyond just positivity, as this would provide a better guide to the appropriate treatment rather than assuming and treating for TB in all of these cases.

